# Context-specific regulation of monocyte surface IL7R expression and soluble receptor secretion by a common autoimmune risk allele

**DOI:** 10.1101/262410

**Authors:** Hussein Al-Mossawi, Nicole Yager, Chelsea Taylor, Evelyn Lau, Sara Danielli, Jelle de Wit, James Gilchrist, Isar Nassiri, Elise A Mahe, Wanseon Lee, Laila Rizvi, Seiko Makino, Jane Cheeseman, Matt Neville, Julian C Knight, Paul Bowness, Benjamin P Fairfax

## Abstract

IL-7 is a key factor in T-cell immunity and *IL7R* polymorphisms are implicated in autoimmune pathogenesis. IL7R mRNA is induced in stimulated monocytes in a genetically determined manner, yet a role for IL7R in monocyte biology remains unexplored. Here we characterize genetic regulation of IL7R at the protein level across multiple cell subsets and conditions in healthy individuals. We find monocyte surface and soluble IL7R (sIL7R) protein are markedly expressed in response to lipopolysaccharide (LPS). We further demonstrate alleles of rs6897932, a non-synonymous IL7R polymorphism associated with susceptibility to Multiple Sclerosis, Ankylosing Spondylitis and Primary Biliary Cirrhosis, form the key determinant of both surface IL7R and sIL7R in the context of inflammation. No effect of this allele was observed in unstimulated monocytes or across lymphoid subsets. Production of sIL7R by monocytes greatly exceeded that of CD4^+^ T-cells, and was strongly associated with both rs6897932 genotype and expression of the splicing factor gene DDX39A. Stimulated monocytes were sensitive to exogenous IL-7, which elicits a defined transcriptional signature. Flow cytometry and single-cell sequencing of synovial fluid derived monocytes from patients with spondyloarthritis showed an enlarged subset of IL7R^+^ monocytes with a unique transcriptional profile that markedly overlaps that induced by IL-7 *in-vitro* and shows similarity to the previously described ‘Mono4’ subset. These data demonstrate disease-associated genetic variants at *IL7R* specifically impact monocyte surface IL7R and sIL7R following innate immune stimulation, suggesting a previously unappreciated key role for monocytes in IL-7 pathway biology and IL7R-associated diseases.

## Introduction

IL-7 is crucial for lymphogenesis and peripheral T-cell homeostasis. Genetic polymorphisms at *IL7R*, the locus encoding the alpha chain of the IL-7 receptor, are associated with predisposition to several autoimmune diseases including Ankylosing Spondylitis (AS), Multiple Sclerosis (MS) and Primary Biliary Cirrhosis^1–3^. A non-synonymous polymorphism within *IL7R* (rs6897932) leads to differential splicing of the 6^th^ exon encoding the trans-membrane domain of the receptor, resulting in expression of a transcript that is translated to form a soluble receptor (sIL7R)^1^. The risk allele is associated with increased circulating levels of sIL7R, and this is thought to prolong the half-life of IL-7^4^. Whilst the cellular source for this receptor has been assumed to be lymphoid, this has not been unambiguously demonstrated, with much of the data derived from cell-lines and mixed cell populations.

An understanding of the cell types and context in which disease risk loci elicit functional activity can provide novel insights into pathogenic mechanisms. Expression quantitative trait locus (eQTL) analysis provides unbiased insights into the genetic determinants of gene expression. In primary monocytes a major eQTL mapping to rs931555 (in linkage disequilibrium (LD) with rs6897932 (r^2^=0.75, D’=0.87) and 53kB 5’ to *IL7R*) is uniquely associated with IL7R expression after chronic exposure to LPS^5^. Despite the lymphoid role of IL7R, eQTL have not been observed in T-cell datasets^6–8^. Given the large effect size of this eQTL on IL7R gene expression after LPS stimulation, and the high LD with a disease-associated polymorphism, we aimed to further characterize this observation at the protein level in immune cell subsets across a large cohort. We performed parallel experiments in isolated monocytes under the same conditions as our previous eQTL analysis, and in mixed cell populations using flow cytometry.

We find monocytes from healthy individuals robustly express surface protein IL7R after exposure to innate immune activators in a manner dependent upon rs6897932 allele. After stimulation, monocytes are the predominant source of sIL7R, again corresponding to rs6897932 allele carriage, with sIL7R levels inversely correlating with the allelic effect on surface receptor. sIL7R protein levels were also correlated with the expression of >1500 transcripts from the same individuals, most significantly being the splicing factor gene *DDX39A*. IL-7 induces a specific transcription profile in IL7R^+^ monocytes, which corresponds to a specific monocyte subset characterized by high expression of *TCF7* and *CCL5*. We demonstrate IL7R^+^ monocytes are comparatively enriched in the synovial fluid of patients with active spondyloarthritis and, using single-cell RNA sequencing, we find synovial IL7R+ monocytes have a discrete expression profile, with similarly marked upregulation of *LTB, TCF7* and *CCL5*. These data demonstrate that genetic variation at *IL7R* impacts IL-7 biology specifically in the setting of inflammation and suggest a key, hitherto unappreciated, myeloid role in the functional mechanism of disease risk variants.

## Results

### Monocyte activation induces expression of surface IL7R

Peripheral blood mononuclear cells (PBMCs) and isolated CD14^+^ monocytes from healthy volunteers were incubated for 24 hours (24h) with or without LPS and surface IL7R was characterized with flow cytometry (Supplementary Figure 1). In both preparations treatment with LPS led to pronounced induction of surface IL7R (Figure 1a-b), demonstrating LPS induction of IL7R mRNA is accompanied by readily detectable surface receptor expression. LPS elicits release of multiple cytokines from monocytes in PBMCs, including IL-4 and IL-6, which alter T-cell surface IL7R^9^. In keeping with this, we observed CD4^+^, CD8^+^ T-cells and CD56^+^ NK cell surface IL7R fall markedly upon stimulation (Figure 1c). We found high inter-individual variation in IL7R response to LPS, particularly in monocytes, and observed little correlation between cell types (Figure 1d), indicative of cell-type specific regulation. Notably, whilst within-cell type correlation of IL7R^+^ between treated and untreated lymphoid cells was high, correlation between percentage IL7R^+^ CD14 in untreated and LPS treated states was weak (CD14^+^ within PBMC culture: r=0.20, p=0.03; purified monocytes: r=0.05, p=0.66) highlighting the cell-type and context-specificity of IL7R response to LPS. To explore the effect of other innate stimuli we exposed monocytes to Pam3CysK4 and Imiquimod, agonists to TLR1:2 and TLR7 respectively. Whilst Pam3CysK4 induced monocyte IL7R expression in all individuals, the response to Imiquimod was variable and non-significant (Figure 1e). Since LPS elicits the release of multiple cytokines, we reasoned that late IL7R induction may relate to magnitude of early (autocrine) cytokine response. To test this, the expression of monocyte IL7R 24h post LPS was correlated with the expression of all genes following two hours (2h) of LPS in 228 individuals^5^. We noted 382/15421 probes (FDR<0.05) where the 2h mRNA expression was correlated with 24h IL7R mRNA expression (Supplementary table 1). Within the top 10 associated genes we found both tumour necrosis factor (*TNF*) and chemokine ligand 5 (*CCL5*) were strongly associated (*TNF:* r=0.34, P adj.=2.9^−4^; CCL5: r=0.36, P adj.=1.1^−4^; Figure 1f). Given the key role for anti-TNF therapies in the treatment of many autoimmune conditions, we tested whether released TNF may drive monocyte IL7R induction in an autocrine manner by adding the anti-TNF monoclonal antibody Infliximab to the media with LPS. This significantly reduced IL7R^+^ monocytes (mean −8.1%; 95% CI: −5.7:-10.5%; P=5.6^−9^, Figure 1g). Finally, we directly tested TNF in a further 78 individuals, incubating cells with TNF for 24h. This showed that, akin to LPS, TNF alone robustly induces the expression of surface IL7R in monocytes (Figure 1h), whilst concomitantly reducing T-cell IL7R surface levels.

**Fig. 1.**
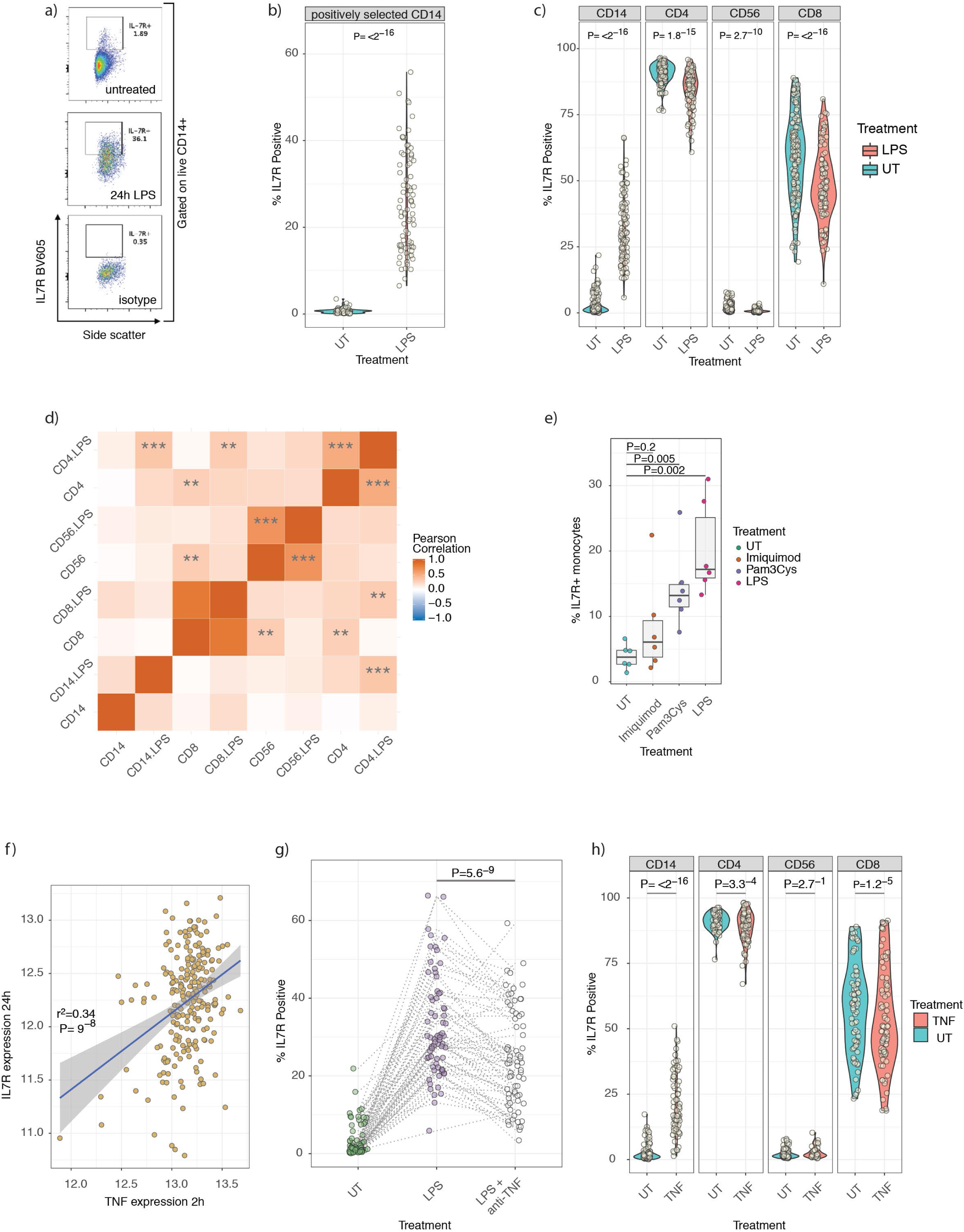
LPS induces profound monocyte IL7R cell surface expression. 1 a) Representative flow cytometry plots from live gated positively selected CD14 monocytes from one individual where cells were incubated for 24h alone (top panel) or in presence of LPS (bottom panels) from same individual 1 b) Violin plot demonstrating significant induction of IL7R^+^ monocytes in across paired cultures of positively selected monocytes (Monocyte IL7R^+^ post LPS-median:23.8, min:6.5, max:55.8, IQR:15.6-34.9%; n=84, paired T-test) 1 c) Comparative effects of LPS on IL7R+ counts across CD14^+^ monocytes, CD4+ and CD8^+^ T-cells and CD56^+^ NK cells from the same PBMC cultures, untreated or with LPS for 24h (PBMC IL7R^+^ post LPS-median:29.9%, min:5.9, max:66.40, IQR:22.3-40.0; n=103, paired T-test) 1 d) Pearson correlation analyses were performed between indicated cells and treatments on number of IL7R^+^ cells. *** P <0.001, ** P<0.01. 1 e) Comparative induction of IL7R^+^ monocytes from 6 individuals with monocytes treated for 24h with either the TLR7 agonist Imiquimod, TLR1/2 agonist Pam3CysK4 or TLR4 agonist LPS (one way ANOVA). 1 f) Expression of TNF at 2h LPS assayed versus expression of IL7R from monocytes from same individuals at 24h LPS. 1 g) Comparative incubation of monocytes alone (n=69), with LPS or with LPS + anti-TNF monoclonal antibody (Infliximab). Incubation with TNF antagonist significantly reduces LPS induced IL7R^+^ monocyte counts (paired t-test). 1 h) Violin plots of responses across cell types of PBMCs treated with TNF alone leads to significant induction of IL7R^+^ monocytes and significantly reduces CD4 and CD8 T-cell IL7R^+^ positivity (n=78, paired T-test).

### rs6897932 specifically regulates monocyte cell surface IL7R levels

In eQTL analysis of stimulated monocytes the most significant association for *IL7R* mRNA expression after exposure to LPS for 24h mapped to rs931555 (P=2.1^−26^), which was also detectable after 2h LPS (Figure 2a). Interrogating these data further, we found rs6897932 associated with *IL7R* expression (P=8.6^−14^) reflecting the degree of LD between these two loci. Conditioning on rs931555 resolves the rs6897932 association (P=0.92, Supplementary Figure 2), whereas controlling for rs6897932 leaves a residual effect of rs931555 (P=7.6^−11^), indicating rs931555 marks the functional haplotype at the mRNA level. After completion of flow-cytometry experiments, the cohort was genome-wide genotyped on the Illumina OmniExpress, with imputation using the UK10K dataset^10^. We then explored the genetic association with surface expression in purified monocytes post LPS stimulation. This demonstrated the most significant local association with surface expression of IL7R was to rs6897932, with no significant association in the unstimulated samples (Figure 2b,2c). In monocytes gated within stimulated PBMC, a significant association with rs6897932 was similarly apparent (Figure 2d). Of note, we were unable to observe a genetic association between surface IL7R and rs6897932 in either CD4^+^ or CD8^+^ T-cells, in the basal (Supplementary Figure 3) or LPS/TNF-stimulated state. We had reasoned that if genetic regulation of IL7R expression in T-cells was context dependent then an alternative stimulant to LPS or TNF may be required. To explore this, we tested the effect of T-cell-specific stimulation with CD3/CD28 ligation in a subset of samples. This reduced T cell IL7R surface levels whilst inducing monocyte surface IL7R without evidence of a genetic association (Supplementary figure 4). Similar to LPS, we found rs6897932 genotype was associated with monocyte surface IL7R after TNF, with the disease risk allele showing lower average surface IL7R (P<0.0007, Figure 3e). These data demonstrate rs6897932 has a previously unrecognized role in regulating the surface expression of monocyte IL7R after exposure to inflammatory mediators LPS and TNF.

**Fig. 2.**
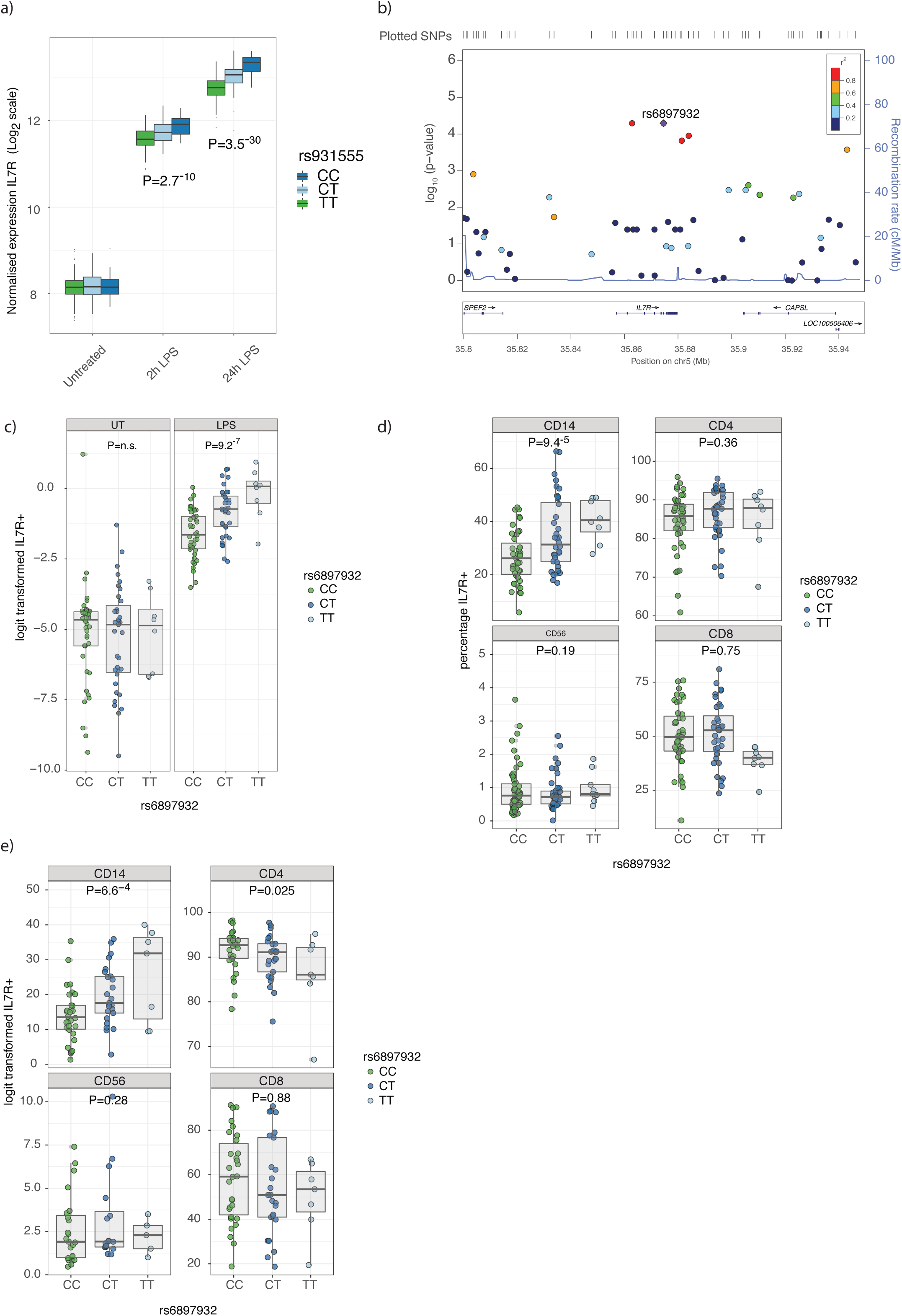
rs6897932 specifically regulates monocyte cell surface IL7R expression. 2 a) Microarray data demonstrating eQTL to IL7R noted at rs931555 after both 2 and 24 hour treatment with LPS (2h LPS: n=261, 24h LPS n=322) 2 b) Local association plot for monocyte surface IL7R after exposure to LPS from positively selected monocytes demonstrates peak association to rs6897932 (n=84) 2 c) Batch corrected log values for surface IL7R from positively selected monocytes demonstrating significant effect of rs6897932 carriage after stimulation in comparison to baseline monocytes (n=84, ANOVA for linear fit model) 2 d) Surface IL7R in PBMCs across cell subsets after exposure to LPS. (n= 87, Note a significant effect of rs6897932 on surface IL7R is only observed in CD14^+^ monocytes, ANOVA for linear fit model) 2 e) As per 2 d) but PBMCs treated with TNF for 24h (n=62)

### Monocyte derived sIL7R is regulated by rs6897932 carriage and associated with DDX39A expression

The reported role of rs6897932 in determining splicing of a soluble form of IL7R in PBMCs^4^ led us to investigate whether stimulated monocytes produce sIL7R and if this is under allelic control. We measured sIL7R from supernatants from randomly selected monocyte samples treated with LPS for 24h from our previous eQTL study (n=161). Notably, we found monocytes secreted large amounts of sIL7R, the yield being robustly influenced by rs6897932 carriage (mean CC allele=3149.5ng/ml, TT allele=916.9ng/ml, P allelic effect=4.7^−15^, Figure 3a). We found only a weak effect of the eQTL locus rs931555 on sIL7R alone (P=0.006), although when controlling for rs6897932 we found rs931555 has an additive effect on sIL7R (Figure 3b), consistent with the upstream eQTL acting independently of rs6897932 to modulate sIL7R.

**Fig. 3.**
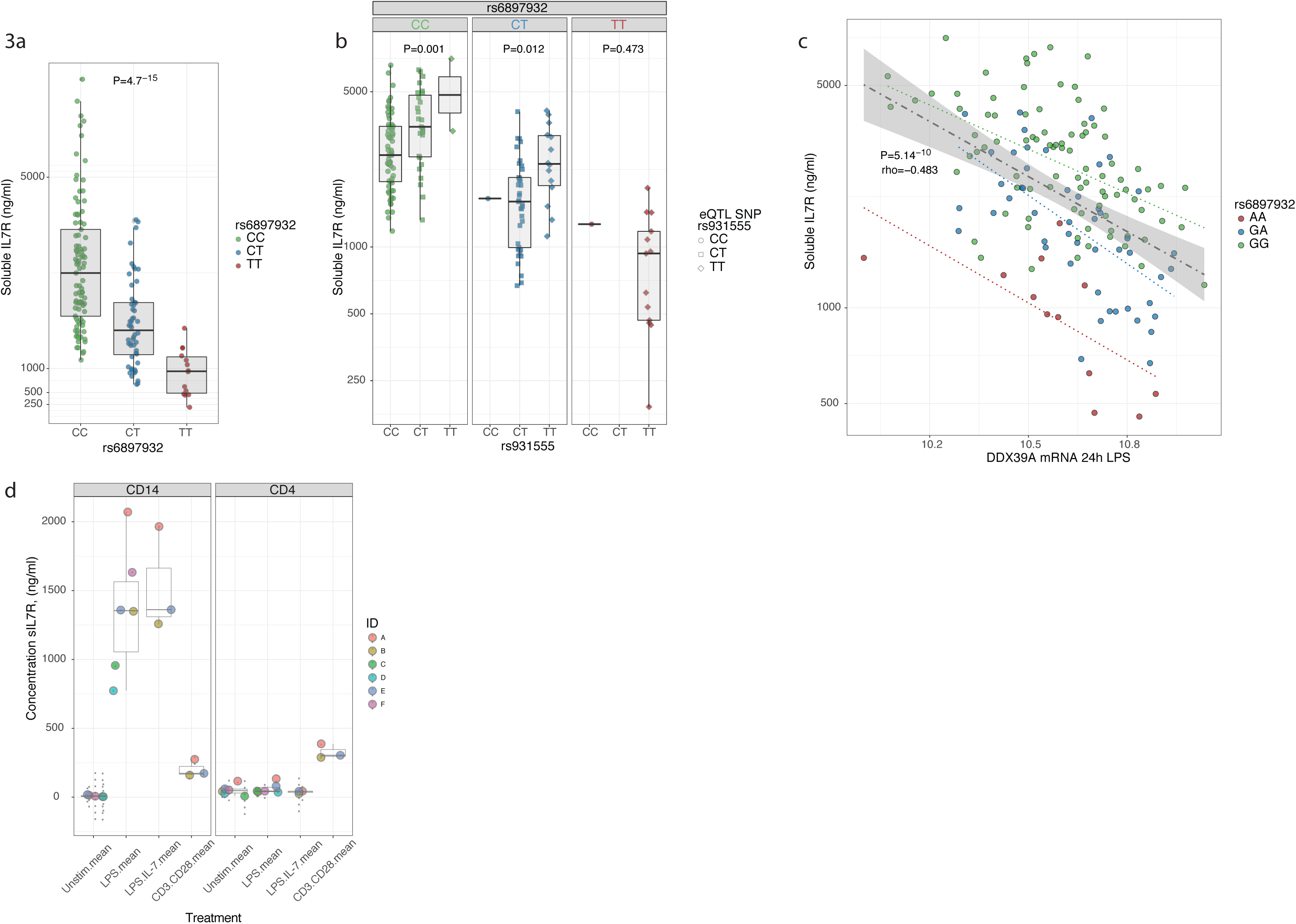
Monocyte-derived soluble IL7R is regulated by rs6897932 carriage and DDX39A expression. 3 a) Elisa of sIL7R performed on supernatants of monocytes treated with LPS for 24h. Samples were randomly chosen from original eQTL dataset and genotypes revealed *post-hoc* – a significant association was observed at rs6897932 (n=161, ANOVA for linear fit model). 3 b) Elisa data from 3 a) showing additional effect of rs931555 genotype on sIL7R (linear model). 3 c) Correlation analysis performed between 15421 microarray probes and soluble IL7R (y-axis) identified DDX39A expression (x-axis) to be significantly anti-correlated with soluble IL7R levels, supporting evidence for its regulation of soluble IL7R. Trend lines indicate correlation per by allele, with rho and P-value for all data (n=161) 3 d) Elisa of sIL7R from isolated primary CD14 and CD4 cells cultured for 24 hours either alone or in the presence of LPS, LPS + IL-7 and CD3/CD28 activating beads. (n=6).

It has been reported that rs2523506, at the 6p21 gene *DDX39B*, is in epistasis to rs6897932 and that allelic combinations of these two loci further increase the risk of MS^11^. *DDX39B* encodes an RNA helicase that forms a component of the spliceosome^12^. Risk polymorphisms in this gene were associated with increased sIL7R in rs6897932 risk allele carriers^11^. In light of these findings, we proceeded to explore this association in our cohort. We did not observe an effect of rs2523506 on rs6897932 regulation of sIL7R in monocytes (Supplementary Figure 5), although this may reflect, even in this relatively large cohort, the low frequency of the minor allele at rs2523506 diminishing power to replicate an epistatic effect. To further understand transcriptional drivers of sIL7R we used pre-existing gene expression data from these samples to explore associations between mRNA expression and sIL7R. In support of the relative independence of soluble protein and total transcript we see no association between total IL7R transcript and sIL7R (Supplementary Figure 6a). We similarly did not observe an association between *DDX39B* expression (at either 2h or 24h of LPS – 3 probes to *DDX39B* (*BAT1*) on the array, all P>0.1) and sILR. However, in strong support of the postulated role of the spliceosome on sIL7R levels, we found a significant correlation between expression of the *DDX39B* paralogue, *DDX39A*, after 24h LPS and sIL7R, with increased expression of *DDX39A* being associated with reduced sIL7R (2h LPS *DDX39A* mRNA vs. sIL7R rho=−0.25, P=0.01; 24h LPS *DDX39A* mRNA vs. sIL7R rho=−0.483,P=5.1^−10^, Figure 3c, Supplementary Table 2). This is in keeping with the directional effect of DDX39B reported in transfected HeLa cells^11^. Total IL7R transcript and *DDX39A* expression were not correlated (Supplementary Figure 6b), supporting a specific role for *DDX39A* in sIL7R splicing as opposed to expression. Finally, we tested the relative production of sIL7R in monocytes versus CD4^+^ T-cells, which show the highest levels of IL7R surface expression (Figure 1c). Whereas neither untreated CD14^+^ or CD4^+^ cells release sIL7R, only CD14^+^ cells release sIL7R in response to LPS, and this was not influenced by IL-7. Conversely, in spite of constitutively expressing far higher levels of IL7R, CD4^+^ T-cells release very little sIL7R, even when stimulated with CD3/CD28 beads. These data demonstrate that, in the context of inflammation, CD14^+^ monocytes form a major source of sIL7R, supporting the importance of the monocyte specificity of the disease associated allele.

### Monocyte surface IL7R is sensitive to IL-7 and stimulation activates multiple transcriptional pathways

To explore a role for IL-7R signalling in monocytes we co-incubated fresh PBMCs with LPS in the presence of recombinant human IL-7 and performed flow cytometry for a subset of individuals. We confirmed previous observations of reduced IL7R expression on T-cells^9^ following IL-7 exposure and similarly found IL-7 markedly reduced monocyte surface IL7R expression, indicating sensitivity of monocyte IL7R to free IL-7 (Figure 4a). To determine the transcriptional consequences of ligation of IL7R in monocytes, monocytes (pre-incubated with LPS to induce IL7R) were exposed to IL-7 for 2h. RNA sequencing was performed on these cells, as well as cells treated with LPS alone, from the same individuals. Pairwise differential expression analysis demonstrated marked transcriptional activity from this short exposure to IL-7 with 3240/16186 transcripts tested being differentially expressed (FDR<0.05, Supplementary table 3 and Figure 4b,c). These included the transcription factor *TCF7*, which is required for immature thymocyte survival^13^, the transcriptional activator *MYB*, the non-coding RNA *MALAT1* and known IL-7 regulated cytokines *LTA* and *LTB* which are crucial for the formation of embryological peripheral lymph nodes^14^. Gene ontology analysis of induced genes was consistent with the known anti-apoptotic effects of IL-7 with significant enrichment of genes involved in translation and ribosomal small subunit biogenesis (Figure 4d).

**Fig. 4.**
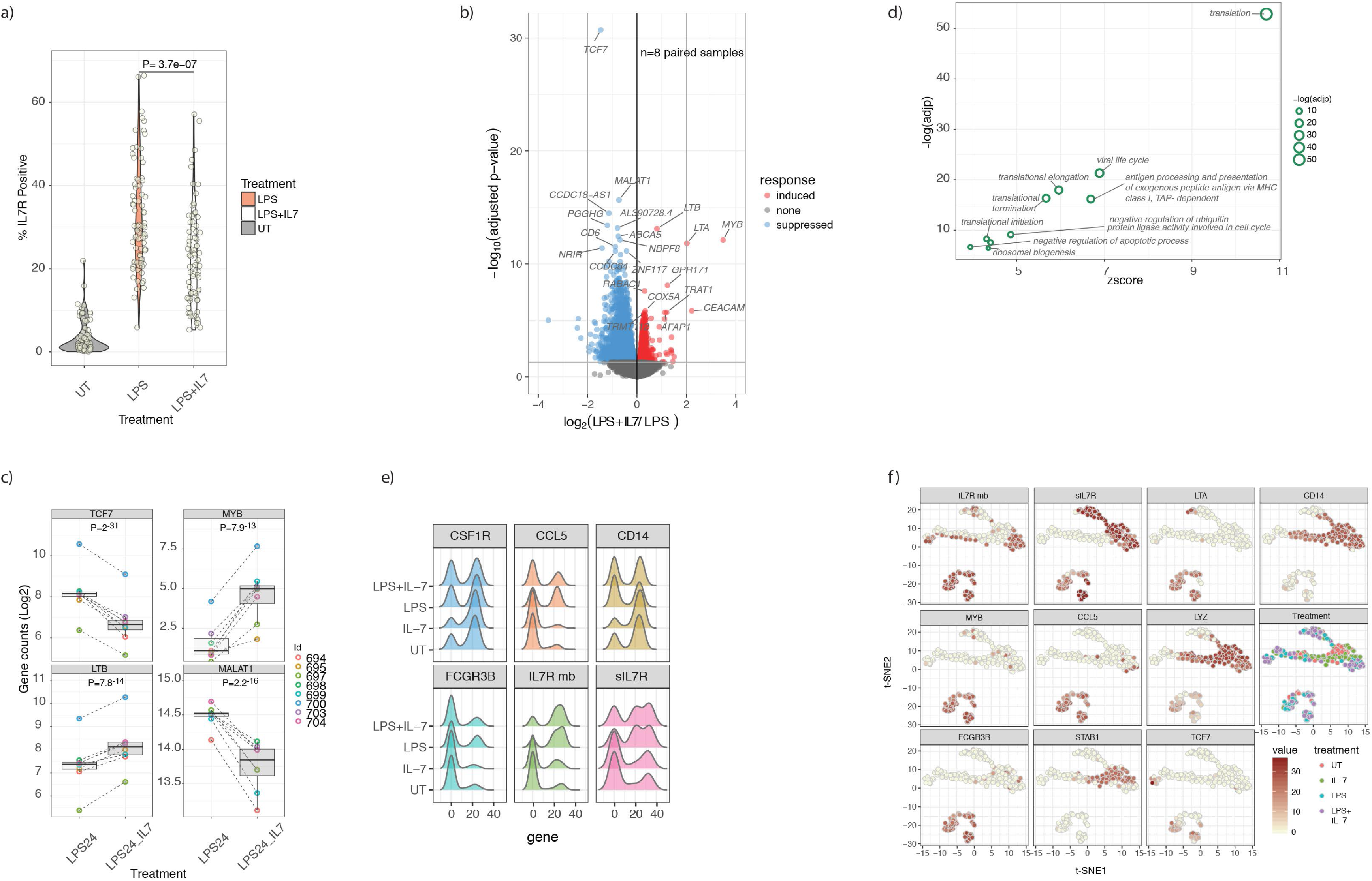
Monocyte surface IL7R is functional and triggers transcriptional activity upon ligation. 4 a) Violin plot demonstrating monocytes from PBMC cultures in the untreated state, after exposure to LPS and after exposure to LPS with recombinant IL-7. IL-7 leads to marked downregulation of IL7R on LPS stimulated monocytes, indicative sensitivity to exogenous cytokine (paired T-test). 4 b) Volcano plot of mRNA from RNAseq experiments of 8 paired monocyte samples either treated with LPS alone for 24h or with LPS with additional IL-7 added for the last 2h of culture. Treatment leads to widespread differential transcript expression with the most significant 20 transcripts labelled (FDR<0.05). 4 c) Example boxplots of genes differentially regulated by recombinant IL-7 in monocytes (linear model) (n=8 subjects). 4 d) Results from gene ontology biological pathway analysis was performed on IL-7 induced genes with all significant pathways labelled. 4 e) Single-cell real-time quantitative PCR (RT-qPCR) data from 3 individuals (192 cells) either untreated or in the presence of IL-7 alone, LPS alone, or LPS with IL-7 added in the last 2h of culture showing expression of *CSF1R, CCL5, CD14, FCGR3B*, membrane-bound IL7R (IL7Rmb) and sIL7R. 4 f) t-SNE plot of single-cell RT-qPCR data showing expression of IL7Rmb, sIL7R, *LTA, CD14, MYB, CCL5, LYZ, FCGR3B, STAB1* and *TCF7* overlaid on the unbiased clustering.

Whilst the bulk sequencing data showed transcriptional sensitivity of monocytes to IL-7 post LPS stimulation, it did not illuminate whether cells expressing IL7R co-express IL-7 responsive genes. To investigate this, we performed single-cell quantitative PCR on monocytes, testing transcripts selected on response to LPS or IL-7 including soluble and full-length IL7R, across the 4 stimulation states (Untreated, IL-7, LPS, LPS+IL-7). This again showed induced expression of both IL7R transcripts by LPS (Figure 4e). T-distributed stochastic neighbour embedding (t-SNE) analysis of these data identified a subset of monocytes strongly positive for IL7R that co-expressed CCL5, a correlate of IL7R expression in the array data, and IL-7-responsive genes including *LTA, MYB* and *TCF7* (Figure 4f).

### IL7R^+^ monocytes form a distinct subset detectable in synovial fluid

Although our data clearly show the ability of monocytes to induce and express surface IL7R in response to stimulation, the physiological and pathological significance of these observations remains uncertain. To explore the relevance of these cells in an inflammatory disease state, we performed pairwise flow cytometry from blood and synovial fluid in 4 patients with spondyloarthritis (2 AS, 2 Psoriatic Arthritis (PsA)). We found that, whereas IL7R^+^ monocytes were infrequent within PBMCs from these individuals, they were readily observed in the synovial fluid from actively inflamed joints, comprising 12-35% of the monocyte population (Figure 5a,b). We used single-cell RNA sequencing of paired blood and synovial fluid from a further three individuals with spondyloarthritis to investigate the CD14 myeloid compartment. Unbiased t-SNE clustering of 1291 synovial monocytes revealed 5 distinct clusters (Figure 5c), with cluster 4 showing high IL7R expression (Figure 5d) relative to the other monocyte clusters. Significantly co-expressed genes defining the IL7R^+^ monocyte cluster included *LTB, CCL5* and *IL-32* (Figure 5e,f, full gene list supplementary table 4). Comparison with the in-vitro stimulated bulk monocytes demonstrated a significant overlap with the IL-7 responsive genes (P=8.5^−4^; OR 2.9; 95% CI: 1.5:5.1), Fisher’s exact test). In the paired PBMC monocytes the same clustering parameters only revealed 3 monocyte clusters with no distinct IL7R^+^ cluster (Supplementary Figure 7a-c). Overall these data confirm that IL7R^+^ monocytes are found in inflamed spondyloarthritis synovial fluid ex-vivo and may represent a unique subset of monocytes based on single-cell gene expression.

**Fig. 5.**
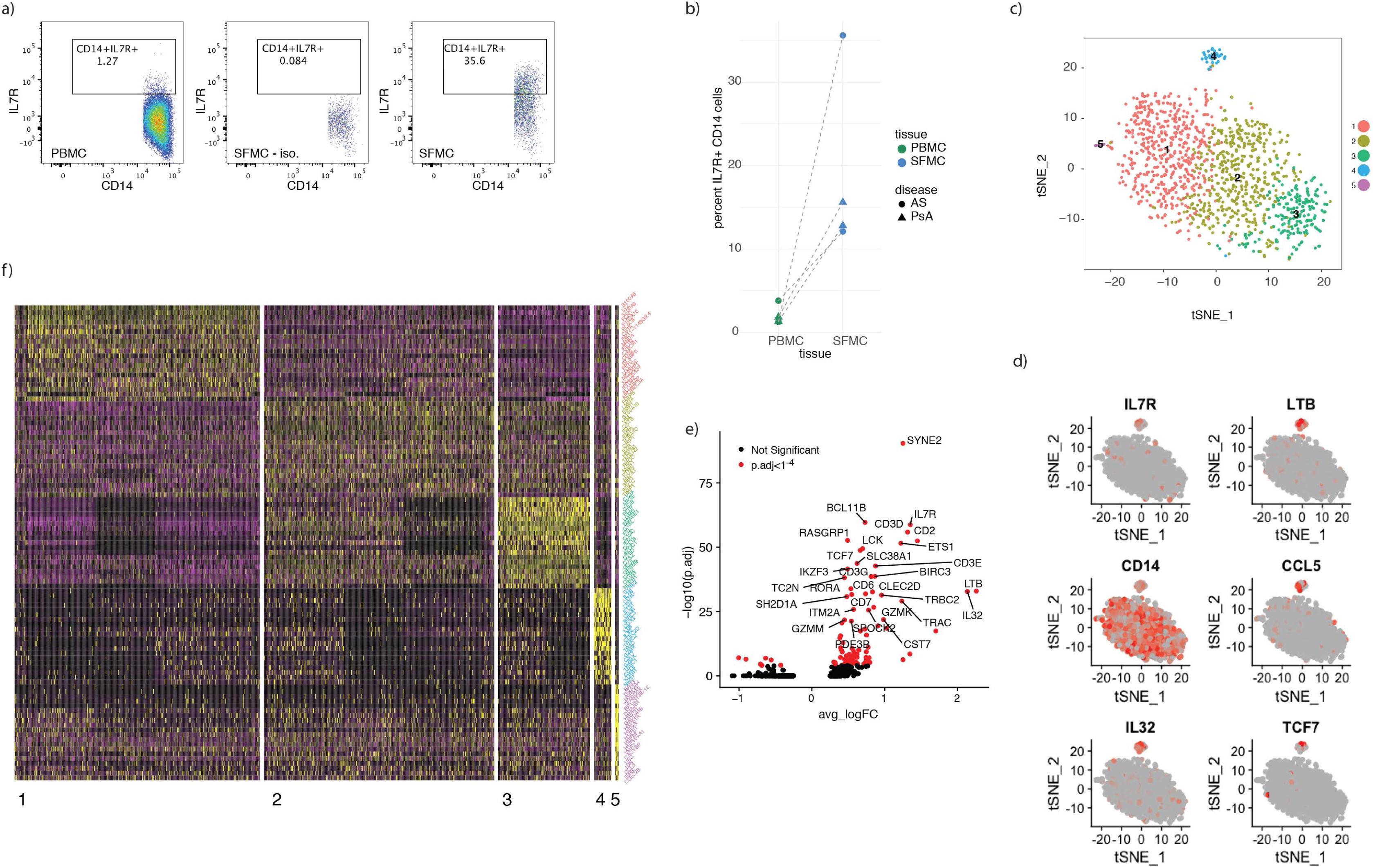
Synovial fluid from spondyloarthritis patients is enriched for IL7R^+^ monocytes which carry a distinct transcriptional signature. 5 a) Representative flow cytometry of IL7R staining of matched PBMC (left) and SFMC (middle isotype control, right) from a spondyloarthritis patient. Gated on CD3^−^CD14^+^. 5 b) Results from 4 patients (2 As and 2 PsA) showing monocyte IL7R staining in matched PBMC and SFMC using flow cytometry. 5 c) t-SNE clustering of monocytes from single-cell RNA sequencing of spondyloarthritis patient (n=3) SFMC showing 5 clusters (n=3). 5 d) Single-cell monocyte expression of IL7R, LTB, *CD14, CCL5, IL32* and *TCF7* overlaid on t-SNE clustering in (c). 5 e) Volcano plot of genes significantly upregulated in the SFMC monocyte cluster 4 compared to all other SFMC monocyte clusters with top 20 genes annotated. 5 f) Heatmap of top 20 genes from the each of the five SFMC monocyte clusters identified in (c).

## Discussion

The degree to which eQTL correspond to subsequent protein quantitative trait loci is variable^15^ and relatively poorly characterised for primary immune cells. The IL7R eQTL in monocytes post LPS is particularly intriguing given the large effect size (r^2^=0.30), linkage with a disease associated locus, and the lack of prior exploration of the role of IL7R in monocytes. Here we show that both surface IL7R and sIL7R are robustly expressed at the protein level in stimulated monocytes and the disease-associated polymorphism rs6897932 plays the predominate role in modulating this. As per the eQTL data, the surface receptor QTL is only observed in the stimulated state and is directionally consistent in monocyte and PBMC cultures, reinforcing the reproducibility of these findings. The anti-TNF monoclonal antibody infliximab reduces LPS induction of IL7R. This both implicates autocrine release of TNF, and suggests that our observations may be of pathogenic and clinical significance, since anti-TNF therapy plays a key role in the disease management of the spondyloarthritides. We further confirm that TNF alone can elicit monocyte IL7R expression. In doing so we again replicate the allelic effect of rs6897932 on monocyte surface IL7R in a separate cohort upon treatment with TNF. Monocytes act as primary secretors of TNF during acute inflammatory events and so this observation of monocyte-specific feedback resulting in expression of IL7R is of particular interest given the accumulating evidence for myeloid role in many inflammatory disease states^16^.

Soluble IL7R is thought to potentiate IL-7 signaling^4^ and likely plays an important role in IL-7 biology and disease associations. We find activated monocytes are the main cellular source of sIL7R in a rs6897932-delineated manner suggesting that, in the context of innate immune activity, monocytes may significantly contribute to the local pool of sIL7R. Although we have limited power to determine epistatic effects, analysis of individuals heterozygous for rs6897932 demonstrates an additional significant allelic effect of rs931555 (in the direction of the previously identified eQTL) on the regulation of sIL7R. It would thus appear that rs6897932 is the primary determinant of sIL7R release from monocytes and is also key in determining surface expression. In keeping with a recently described role for the spliceosome component DDX39B in IL7R splicing^11^, we observed a highly significant correlation between expression of the DDX39B paralogue gene, DDX39A, and sIL7R levels, with increasing expression of DDX39A associated with reduced sIL7R. The absence of association to DDX39B may reflect cell type specific mechanisms or potentially a divergence from the steady state induced by stimulation.

We did not observe an effect of rs6897932 on surface expression of IL7R in either CD4^+^ or CD8^+^ T-cells in resting or stimulated states. Given we did not measure T-cell derived soluble IL7R we cannot exclude an allelic effect of rs6897932 on this, but it is notable that stimulated monocyte production of sIL7R is several fold greater than that of stimulated T cells. Interestingly, previous associations of this allele on sIL7R were from mixed cell populations^4^. Furthermore, there is an absence of an eQTL to IL7R in the GTEX data at rs6897932 and an eQTL is not observed in CD4^+^ T-cells^6^. Whereas sIL7R levels are consistently demonstrated to be raised in autoimmune conditions – notably MS^4^, specific attempts to assay sIL7R from primary T cells have not demonstrated an allelic association with rs6897932^17–19^. Our data is thus in keeping with a predominant effect of this allele on both transcript and surface protein expression in activated monocytes, and implicates monocytes as a key contributory cell type to the disease association.

In addition to the role of monocyte sIL7R in modulating inflammation, the direct ability of monocytes to respond to IL-7 via the membrane bound receptor is intriguing. In particular, since circulating IL-7 levels are thought to be regulated by consumption^20^, monocytes may be acting as an IL-7 sink to limit the amount of IL-7 available for lymphocyte consumption. Hence, the higher monocyte surface IL7R expression observed with the protective rs6897932 allele would be in keeping with the low CD8^+^ T-cell count previously reported in carriers of the same allele^2^, an observation reinforced by analysis of UK biobank datasets where rs6897932 is associated with lymphocyte percentage^21^. Moreover, the direct modulation of genes by IL-7 observed in-vitro and the significant overlap with the *ex-vivo* single-cell data would suggest that IL-7R+ monocytes may represent a unique subset of monocytes with a hitherto unknown role in inflammation biology. Of particular interest is the observation that IL-7R^+^ monocytes express a number of genes previously thought to be T-cell specific such as *CD2* and *LCK*. Notably, such a subset of monocytes, labelled ‘Mono4’, which similarly over-expresses *CCL5, IL32* and *IKZF3*, was recently reported from a flow-sorted smart-seq2 data-set^22^. Our data places these cells at the inflamed spondyloarthritis joint and suggests this subset may be playing a role in inflammation. Whether such cells have separate immunomodulatory potential warrants further investigation.

IL7R can also dimerize with the thymic stromal lymphopoietin receptor^23,24^, however monocyte expression of this is very low and it is not induced by LPS^5^, thus we focused in this study to investigate the role of IL-7 on monocyte biology. Whilst we demonstrated TNF induces IL7R expression, the degree of this response tended to be less than to that of LPS. This was unlikely to be a dosage effect, as higher concentrations of TNF led to cell death (data not shown), but instead suggests a role for other co-released cytokines or the TLR4 pathway directly. Finally, whilst we did not observe an effect of this allele on other cell subsets, our analysis was of crude CD4^+^ and CD8^+^ fractions, and thus allelic modulation of surface IL7R in smaller subsets cannot be discounted. Notably, our proposed mechanism with the disease associated allele increasing monocyte-derived sIL7R in inflammation would likely act via its known effect on potentiating the bioavailability of IL-7^4^. Importantly, this model is in keeping with a role for IL-7 in T-cell driven pathogenesis^25^ and is compatible with the genetics, where large datasets are unable to demonstrate a direct effect of this allele on T-cell expression.

In summary, our study provides unequivocal evidence that the disease associated polymorphism rs6897932 is associated with monocyte specific modulation of surface IL7R and sIL7R in the context of inflammation. We further demonstrate that IL7R-expressing monocytes are responsive to IL-7 with a distinct expression profile that significantly overlaps ex-vivo IL7R^+^ monocytes from the joints of patients with spondyloarthropathies. These observations highlight a previously unrecognized role for monocytes in IL-7 biology and may have implications in development of new therapies for autoimmune diseases.

## Supplementary figures

S1 Gating strategy for PBMC IL7R staining

S2 Association plots corrected for rs6897932 and rs931555

S3 Surface IL7R by rs6897932 allele in untreated PBMC cultures

S4 Induction of monocyte IL7R^+^ by T-cell specific stimulation with CD3.CD28 beads

S5 Exploration of epistatic effect of rs2523506 and rs6897932

S6 Relationship between IL7R transcript, soluble receptor and DDX39A

S7 Monocyte single-cell RNA seq data from spondyloarthritis PBMC

## Supplementary tables

T1 Genes where expression at 2h LPS is correlated with 24h LPS IL7R

T2 Genes correlated with soluble IL7R at 24h LPS

T3 Genes differentially regulated by IL-7

T4 Genes differentially expressed in IL7R cluster from SFMC single-cell RNA seq

T5 Antibody clones for flow cytometry

## Materials and Methods

### Study participants

Peripheral blood was obtained from genotyped individuals recruited via the Oxford biobank (www.oxfordbiobank.org.uk) with full ethical approval (REC: 06/Q1605/55) and written informed consent. Genotype was blinded at the time of study and only revealed at the end of recruitment. Peripheral blood and synovial fluid samples from were recruited with informed consent from patients with inflammatory arthritis attending the Oxford University Hospitals NHS Foundation Trust (Ethics reference number 06/Q1606/139).

### Cell isolation and stimulation

Whole blood was collected into EDTA tubes (BD vacutainer system) and peripheral blood mononuclear cells were obtained by density centrifugation (Ficoll Paque). CD14 cell isolation was carried out by positive selection (Miltenyi) according to the manufacturer’s instructions. Cells were rested overnight (16h) at 37°C, 5% CO_2_ in 5ml non-adherent polypropylene cell-culture tubes (BD Biosciences) prior to stimulation assays. Cells were stimulated for 24 hours in RPMI1640 media with 20% fetal calf serum in the presence of 20ng/ml LPS (Invivogen), 100 ng/ml pam 3cysk4 (Invivogen), 100ng/ml imiquimod (Invivogen), 10ng/ml TNF (R&D systems), 10ng/ml IL-7 (Peprotech) and CD2/3/28 beads (Miltenyi) at a ratio of 1 bead to 2 cells. In all experiments an unstimulated incubator control was included. TNF blockade was achieved with 5μg/ml of infliximab (Remicade, Janssen).

### Flow cytometry

Staining antibodies and dye clones, dilutions and manufacturer shown in supplementary table 1. Cells were stained in phosphate buffered saline containing 1% fetal calf serum on ice and in the dark for 20 minutes, then fixed in 1.6% paraformaldehyde. All samples included fixable amine reactive viability dye and isotype control for IL-7R. Flow cytometry was performed on a BD Fortessa calibrated daily with calibration and tracking beads from BD Biosciences. Data was analysed using FlowJo software (Treestar®).

### ELISA

Cell supernatants from stimulated cells were collected and sIL7R quantified as previously described^11^. Briefly, 96-well plates were coated overnight with 1 μg/ml mouse anti-human CD127 mAb (R&D Systems). Plates were blocked with 5% BSA for 1 hour, washed, and cell supernatants added for 2 hours. Bound sIL7R was detected with 12.5 ng/ml biotinylated goat anti-human CD127 polyclonal Ab (R&D Systems) for 1 h, followed by a 30 min incubation with streptavidin-HRP and a 15 min incubation with TMB peroxidase substrate (Thermo Fisher). The reaction was stopped with H_2_SO_4_ and plates read at 450 nm. A standard curve was generated using recombinant human CD127 Fc chimera protein (R&D Systems).

### RNA extraction

The AllPrep DNA/RNA/miRNA kit (Qiagen) was used for RNA extraction. Cells were spun down and re-suspended in 350μl of RLTplus buffer and transferred to 2ml tubes. Samples were then stored at −80°C for batched RNA extraction. Homogenization of the sample was carried out using the QIAshredder (Qiagen). DNase I was used during the extraction protocol to minimise DNA contamination. RNA was eluted into 35μl of RNase-free water. The RNA amount was quantified by qubit and the RNA samples stored at − 80°C for storage until ready for sequencing.

### Bulk RNA sequencing

RNA sequencing was carried out at the Wellcome Trust Centre for Human Genetics core facility. RNA underwent quality control testing using a bioanalyser (RNA 6000 Nano kit, Agilent) followed by cDNA library preparation. Paired end sequencing was performed at 100 base pairs on each side of the DNA fragment on the HiSeq 4000 platform. 47-69 million reads were sequenced per sample (mean=56.5 million) with samples multiplexed to 8 samples per lane.

### Analytical Methods for bulk RNA sequencing

Bioinformatic analysis of RNA sequencing samples was carried out using validated packages in R. Reads were mapped to human genome reference sequence GRCh38 with HISAT^26^. Gene counts were retrieved using HTseq-count^27^ and the Ensembl gene annotation. DESeq2^28^ was used for differential expression analysis using a pairwise model where individual was coded for. Pathway and network analysis was performed using the R package XGR^29^. All statistical analysis was performed using base R. Batch correction was peformed by incorporating batch within linear regression of logit transformed cell counts. Plots were created using the ggplot2 package^30^ and association plots using LocusZoom^31^

### Cell preparation for single-cell RNA-seq

Blood and synovial fluid was collected from patients with peripheral spondyloarthritis and mononuclear cells were isolated immediately by density centrifugation. Cells were then counted and loaded onto a 10X Genomics Chromium machine within 4 hours of collection from the patients. Cells were captured and cDNA preparation performed according to the Single Cell 3’ Protocol recommended by the manufacturer^32^. libraries were Sequenced on an Illumina HiSeq 4000 to achieve 75 bp reads.

### Single-cell real-time quantitative PCR

CD14^+^ monocytes were collected from healthy donors and cultured as above in the presence or absence of 20ng LPS (InvivoGen) for 24 hours (+/−10ng/ml IL-7 (Peprotech)) for 2 hours before single-cell fluorescence activated cell sorting into a 96-well plate. Cells were lysed and reverse transcribed prior to pre-amplification using Superscript III with Platinum taq (Thermo Fisher Scientific) and Superase-IN RNase inhibitor (Ambion). Cells were pre-amplified to amplify any lowly expressed transcripts using Taqman gene expression assays (Thermo Fisher Scientific) and diluted in TE buffer. Single-cell qPCR was performed using Taqman Universal PCR Mastermix (Thermo Fisher Scientific), 192:24 integrated fluidic circuits (Fluidigm) and the Biomark HD system following the manufacturer’s protocol. Cell-free wells and reverse transcriptase-free controls were included as negative and genomic DNA controls as well as a positive control well containing 100 cells. Data was analysed using Real-Time PCR Analysis software from Fluidigm.

### Single-cell RNA-seq analysis

Sequencing reads were mapped with **CellRanger V2.1.0 GRCh38 (ENSEMBL annotation**). Quality control, filtering and clustering analysis was carried out with the R package Seurat version 2.3.1. To exclude low quality cells, cells with fewer than 500 genes excluded. Likely doublets were removed by filtering out cells with greater than 2000 genes or 10,000 UMIs. All cells with a mitochondrial fraction greater than 7.5% were also excluded. 8219 PBMC cells and 6668 SFMC cells passed the filtering. Library-size normalization was performed on the UMI-collapsed gene expression values for each cell barcode by scaling by the total number of transcripts and multiplying by 10,000. The data was then natural-log transformed using log1p.

PBMC and SFMCs were clustered separately. 9712 (PBMC) and 9409 (SFMC) genes with high variance we selected using the FindVariableGenes function with log-mean expression values between 0.0125 and 4 and dispersion (variance/mean) between 0.25 and 30. The dimensionality of the data was reduced using principle component analysis and 15 principle components (PCs) were identified for downstream analysis. The Louvain algorithm^33^ for modularity-driven clustering, based on a cell–cell distance matrix constructed on the defined PCs was then used. This was implemented using the FindClusters function in Seurat with a resolution of 0.6 to identify 11 distinct clusters of cells in both PBMC and SFMC.

Cluster annotation was assigned using canonical expression markers for key cell types such as CD14 for monocytes. The subset function was then used to extract the raw gene expression matrix of the monocyte cluster for further analysis. The monocyte specific analysis involved re-clustering based 15 principle components and a resolution of 0.6. For PBMC 3 monocyte clusters were identified while in the SFMC compartment 5 SFMC clusters were identified. In the SFMC compartment, the FindAllMarkers function was used to identify the top 20 genes in the SFMC monocyte clusters in order to generate the heatmap. For all single-cell differential expression tests, a non-parametric Wilcoxon rank sum test was used.

## Author contributions

HAM & BPF conceived the project; cell purification and stimulation was performed by EL, CT, BPF, SM and SD; flow experiments were performed by HAM with assistance from JW & SD; samples were procured using the Oxford Biobank organized by JG, JC & MN; NY, EM & LR performed ELISA analysis, BPF, HAM, WL & IN performed statistical and bioinformatic analysis. BPF drafted the manuscript and figures with subsequent contributions from all authors. BPF, PB & JCK jointly supervised the project.

## Acknowledgements

We thank the volunteers from the Oxford Biobank (www.oxfordbiobank.org.uk) for their participation and the NIHR Oxford Biomedical Research Centre which supported the recalling process of the volunteers. The views expressed are those of the author(s) and not necessarily those of the NHS, the NIHR or the Department of Health.

## Funding

This work was supported by a Wellcome Intermediate Clinical Fellowship (201488/Z/16/Z to B.P.F.) and The Academy of Medical Sciences Starter Grant for Clinical Lecturers (B.P.F.); Wellcome Investigator Award (204969/Z/16/Z to J.C.K.), Wellcome Grant (090532/Z/09/Z to core facilities Wellcome Centre for Human Genetics; Arthritis Research UK (20773 to J.C.K.) and the National Institute for Health Research (NIHR) Oxford Biomedical Research Centre (BRC) (SD and PB). H.A.M. was supported by Wellcome Studentship (102288/Z/13/Z). The views expressed are those of the author(s) and not necessarily those of the NHS, the NIHR or the Department of Health.

